# Computational modelling reveals slower safety learning and threat extinction are associated with higher anxiety severity in remote fear conditioning

**DOI:** 10.1101/2025.02.18.638846

**Authors:** Tim Kerr, Kirstin Purves, Thomas McGregor, Michelle G. Craske, Tom Barry, Kathryn J. Lester, Elena Constantinou, Michael Sun, Oliver J. Robinson, Thalia C. Eley

## Abstract

Anxiety disorders are are chronic, pervasive, and debilitating; characterised by a persistent or exaggerated response to distal or abstract threats. Impaired threat discrimination (distinguishing safe from threatening stimuli) and impaired threat extinction (learning a once threatening stimulus is now safe), are known risk factors in the development and persistence of anxiety disorders. These effects can be experimentally elicited through fear conditioning. First, repeated trials of paired aversive and neutral stimuli are delivered during a fear acquisition phase, followed by repeated trials with no aversive stimuli in a fear extinction phase. The effects are typically measured through comparison of end-phase data points, or simple descriptive or statistical models. Computational modelling, by contrast, can offer a hypothesis-driven, trial-by-trial mechanistic account of fear conditioning. This unmasks within subject task variance by estimating the rate of threat learning, safety learning, and threat extinction, examining individual differences in the cognitive mechanisms behind anxiety. A normative sample (n = 145) underwent a differential fear conditioning task on a bespoke smartphone app, in addition to completing an anxiety severity measure (GAD-7). Computational models fitted to task data estimated learning rates. Whilst the threat learning rate showed no association, the threat extinction and safety learning rates showed small negative associations with anxiety severity (r = −0.218, p = 0.008 & r = −0.214, p = 0.01 respectively). These findings are in keeping with prior studies using traditional analytical approaches, and indicate that anxious individuals are not quicker to develop fear of a stimulus, but take more time than their non-anxious counterparts to learn that a stimulus is safe. This study strengthens the evidence for impairments in fear extinction in those with anxiety, and the importance of learning rates as an index of anxiety severity, a previously hidden cognitive mechanism underlying anxiety persistence.

## Introduction

### Anxiety disorders

Anxiety is the state, comprising subjective and physiological responses combined with behavioural urges, elicited by non-imminent threats (LeDoux & Pine, 2016). Through an associative learning mechanism, anxiety normally adapts to protect us from novel threats, whilst facilitating exploration when threats are no longer relevant. In pathological anxiety, the subjective, physiological, or behavioural responses are excessive in relation to the level of threat, or persist long after the initial event (Rosen & Schulkin, 1998). The symptoms generated by pathological anxiety interfere with the activities of normal life, with distinct symptom clusters pointing to a diagnosis of one of several anxiety disorders (National Institute for Health and Care Excellence [NICE], 2014). Given their lifetime prevalence of one third (Bandelow & Michaelis, 2015), a typical onset in the second or third decade of life (Kessler et al., 2005; Kessler et al., 2012), and treatment resistance (Bystritsky, 2006), anxiety disorders impart a high societal cost.

### Fear conditioning

Fear conditioning is a commonly used aversive learning paradigm, capable of revealing several facets of threat processing - threat learning, safety learning, and threat extinction (Lonsdorf et al., 2017a). Typically beginning with an *acquisition* phase, threat acquisition is invoked when an initially neutral cue is repeatedly paired with an aversive stimulus. At each presentation of the cue, the conditional, anticipatory threat responding can be measured via physiological, behavioural, or self-reported changes. Anxiety patients have been shown to exhibit greater physiological responding to conditional threat cues (threat cue, CS+) (Lissek et al., 2005), but this effect may be small, and there was little evidence to support the effect in a second meta-analysis (Duits et al., 2015). Interestingly, depressive patients are shown to exhibit impaired fear acquisition, suggesting separate mechanisms may underpin these two often comorbid diagnoses (Luo et al., 2024; Otto et al., 2014; Wurst et al., 2021).

Safety learning is invoked through concurrent control trials included within the acquisition phase, where a different neutral cue is presented with no aversive pairing (safety cue, CS-). The strength of responding to the CS+ and CS-by the end of acquisition training can be compared to establish threat discrimination. Reduced threat discrimination, the ability to distinguish threatening from safe cues, is widely seen in both children and adults with anxiety disorders; predominantly a result of increased threat responding to the safe cue (Craske et al., 2008; Duits et al., 2015; Waters et al., 2009).

During *extinction* the CS+ is repeatedly delivered without an aversive pairing, lessening the strength of conditional responding. However, impaired threat extinction is often seen in anxiety disorder patients, and is likely a separate learning process from threat acquisition (Duits et al., 2015). There is some evidence to suggest this process is also altered in depression (Dibbets et al., 2015; Wurst et al., 2021).

Fear conditioning paradigms allow the investigation of these distinct learning processes and their role in pathological anxiety (Yamamori & Robinson, 2023). Typically, fear conditioning is performed in the laboratory, however, there is an increasing desire to expand both sample size and diversity beyond the confines of laboratory-based procedures (Gillan & Rutledge, 2021; Ney et al., 2023). Our Fear Learning and Anxiety Response (FLARe) smartphone app enables the remote collection of fear conditioning data (McGregor et al., 2022). Previous research using FLARe demonstrates impaired threat discrimination in anxious subjects, in line with the prior meta-analytic studies of fear conditioning (McGregor et al., 2021).

### Computational modelling

Descriptive measures of fear conditioning data, such as comparing the difference in average responding to CS+ and CS-across a whole phase (threat discrimination), only highlight the presence of an effect, rather than the mechanism subtending the effect. Standard linear statistical models go further identifying probabilistic analogues of the mechanism. Further, both approaches suffer non-optimal fitting, resulting in analytic flexibility within the literature (Lonsdorf et al., 2022; Ney et al., 2018).

In a hypothesis-driven manner, generative computational modelling tests different proposed learning mechanisms against one another, and establishes which mechanism offers the best explanation of the observed data (Friston et al., 2017). The individually fitted free parameters underpinning the proposed mechanisms, such as the learning rates of threat learning, safety learning, and threat extinction learning, can be used to test associations with anxiety severity (Yamamori & Robinson, 2023). Individual differences in these rates, and their associations with anxiety severity, have been demonstrated in both remotely delivered aversive learning tasks (Pike & Robinson, 2022), and laboratory-based fear conditioning tasks (Abend et al., 2022; Gershman & Hartley, 2015; Tzovara et al., 2018). Interestingly, there is some evidence that patterns in learning rates differ between anxiety and depression (Cavanagh et al., 2019), and that computational approaches may be suited to exploring such differences (Huys et al., 2016). However, this modelling approach has yet to be deployed in a remote study of fear conditioning.

### Summary

This study modelled individual differences in fear conditioning learning rates, using data from a smartphone delivered paradigm, and examined their associations with anxiety severity. In line with the existing literature, we pre-registered several hypotheses to test these associations - that threat learning rate would positively correlate with anxiety severity, whereas safety learning, and threat extinction learning rates, would show negative associations with anxiety severity.

## Methods

### Sample

In total, n = 235 participants were tested in this study, pooled from three prior studies. Some were pooled from two of our FLARe app validation studies (n = 47 and n = 68, from studies one and two described in Table 1) (Purves et al., 2019). A later, unpublished validation study recruited n = 120 participants (study three in Table 1).

**Table 1.** Demographic Characteristics.

*Demographic Characteristics*
| Exclusion Status | Study | Demographic and Clinical Measures |  |  |  |  |
| --- | --- | --- | --- | --- | --- | --- |
|  |  | N | Sex = Female | Age | GAD | PHQ8 |
| Pre-Exclusion | Study 1 | 47 | 28 (59.6%) | 22.8 (1.5) | 4.7 (4) | 5.1 (4.3) |
|  | Study 2 | 68 | 43 (64.2%) | 23.1 (1.7) | 4.7 (4.8) | 5.4 (5.5) |
|  | Study 3 | 120 | 83 (69.2%) | 22.8 (1.7) | 6.5 (5.3) | 6.3 (5.1) |
|  | Combined | 235 | 154 (65.8%) | 22.9 (1.7) | 5.6 (4.9) | 5.8 (5.1) |
|  | *Between Group Comparison* |  | H=1.49, p=0.476 | H=1.5, p=0.472 | **H=7.17, p=0.028** | H=3.68, p=0.159 |
| Post-Exclusion | Study 1 | 35 | 21 (60%) | 22.8 (1.5) | 4.9 (3.6) | 5 (3.9) |
|  | Study 2 | 37 | 24 (64.9%) | 23.1 (1.7) | 6.3 (5.3) | 7.2 (5.8) |
|  | Study 3 | 73 | 51 (69.9%) | 22.8 (1.6) | 6.3 (5.3) | 5.8 (4.5) |
|  | Combined | 145 | 96 (66.2%) | 22.8 (1.6) | 6 (5) | 6 (4.8) |
|  | *Between Group Comparison* |  | H=1.06, p=0.588 | H=0.7, p=0.705 | H=0.76, p=0.685 | H=2.31, p=0.316 |
| *Pre-Post Exclusion Comparison* |  |  | W=16898, p=0.938 | W=16061, p=0.846 | W=15402, p=0.363 | W=15589.5, p=0.47 |
| Post-Strict Exclusion | Study 1 | 0 | - | - | - | - |
|  | Study 2 | 29 | 18 (62.1%) | 22.9 (1.6) | 6.2 (5.3) | 7.2 (6.3) |
|  | Study 3 | 59 | 43 (72.9%) | 22.7 (1.6) | 6.7 (5.6) | 6.1 (4.6) |
|  | Combined | 88 | 61 (69.3%) | 22.8 (1.6) | 6.6 (5.4) | 6.4 (5.2) |
|  | *Between Group Comparison* |  | H=1.06, p=0.304 | H=0.16, p=0.686 | H=0.08, p=0.776 | H=0.13, p=0.715 |
| *Post – Post Strict Exclusion Comparison* |  |  | W=6181.5, p=0.625 | W=6304, p=0.745 | W=6060, p=0.52 | W=6153.5, p=0.649 |

The additional participants comprising study three were recruited via the King’s College London trial recruitment portal. Participants responded to an advert and were followed up with instructions from the research group if they met the inclusion criteria - adults between the ages of 21 and 26, with no documented neurological, cardiac, or psychiatric history, and owning a smartphone. Participants were paid £15 if they completed the study. The study was approved by the Kings College London Psychiatry, Nursing and Midwifery Research Ethics Subcommittee (reference HR15/16-2349).

### Task

A fear conditioning paradigm was delivered remotely, via the Fear Learning and Anxiety Response (FLARe) application, which was downloaded and installed onto participants’ smartphone devices. Participants followed instructions relating to the set-up of the experiment, asking them to connect headphones, set their device volume to maximum, and complete the subsequent experiment alone in a quiet room. Task-specific instructions were then delivered.

First, participants were presented with a sequence of conditional stimuli (CS) on screen. These were two differentially sized coloured circles. One circle was usually reinforced (CS+) with the unconditional stimulus (US), an unpleasantly loud noise played through headphones. The other circle was always non-reinforced (CS-). The stimuli serving as CS+ and CS-were counterbalanced between participants. At each presentation of CS, participants were asked to rate the certainty with which they expected to hear the US.

The fear acquisition phase consisted of 24 trials in a pseudo-randomised order, twelve trials each of CS+ and CS-. US occurred on 75% of the CS+ trials, i.e. nine of the twelve CS+ trials. Each trial was eight seconds in duration, participants entered their US expectancy rating after three seconds. Each trial was separated by an inter-trial interval of either one, two, or three seconds duration.

Following a ten minute break, in which participants completed questionnaires, participants completed the fear extinction phase, where 36 unreinforced trials were presented to participants, 18 each of CS+ and CS-. At no point was a US delivered. Participants entered their US expectancy rating during each trial, as per the acquisition phase.

Compliance with the task procedure was assessed through a post-experiment questionnaire, where participants were asked if they removed their headphones, or restarted the application at any point (with the caveat that participants would still receive remuneration if they answered yes to these questions). Participants were also asked how unpleasant they found the US on an integer scale of 1-10, and whether they established contingency awareness (i.e. awareness of the pairing between the US and the CS+). Finally, objective data, such as headphone volume manipulation, and application restarts, was automatically gathered by the software (McGregor et al., 2022).

### Measures

#### US expectancy ratings

US expectancy ratings were entered via an ordinal, discrete scale from one to nine (US expectancy rating, nine representing maximal certainty of US, one representing minimal certainty of US).

#### GAD-7

During the ten minute break, participants completed the Generalised Anxiety Disorder Assessment (GAD-7) (Spitzer et al., 2006). This contains seven questions pertaining to the severity of the symptom criteria for Generalised Anxiety Disorder within the DSM-4. Each question is scored on an ordinal categorical scale (Likert scale), ranging from “Not at all” to “Nearly every day”. This scale is converted into an integer score in the interval (0,3), with a maximum total score of 21.

#### PHQ-8

Participants also completed a Patient Health Questionnaire-8 (PHQ-8) measure, a modified version of the PHQ-9, with the question asking about suicidal thoughts removed, given the inability to safety-net a remotely delivered measure. Similarly to the GAD-7 measure, the PHQ-8 contains eight questions probing the symptoms of depression, per DSM-5 criteria. This is scored on an ordinal categorical Likert scale, ranging from “Not at all” to “Nearly every day”. This scale is converted into an integer score in the interval (0,3), with a maximum total score of 24.

### Exclusions

In line with previous studies using the FLARe app, participants who removed their headphones, reduced their headphone volume to below 80% of the maximum, restarted the application during either phase, or rated the unpleasantness of the US as less than five (out of ten) were excluded from the analysis.

### Statistical analysis

Conventional descriptive measures of fear conditioning were initially applied to the data, to allow comparison with the novel computational modelling approaches. These were 1) end phase CS discrimination - the difference between the US expectancy rating for CS+ and CS- at the final trial of either phase, and 2) the mean of the US expectancy ratings for each CS and phase. These offered a subject level measure of threat sensitivity, and a crude account of the trial data preceding end phase scores. Both measures were correlated with GAD-7 scores, to examine associations between fear conditioning measures, and anxiety severity. Non-parametric Wilcoxon Signed Rank tests were used to assess CS discrimination, given the bounded and therefore non-normal distributions of expectancy rating data. Likewise, given the zero-inflated, ordinal nature of the GAD-7 score, the non-parametric Spearmans Rho correlation was used to estimate associations.

### Pre-registration

The study was pre-registered on the Open Science Framework (OSF) prior to initial computational modelling analysis (https://doi.org/10.17605/OSF.IO/QKBXP). In the process of model fitting to real data, the model specifications deviated from this plan due to computational intractability of some of the proposed models. This did not however impact the hypotheses registered, which could still be tested via the remaining models.

### Computational modelling

Twenty eight Rescorla Wagner associative learning models were fitted to the data, with estimated individual learning rates weighting the trial-by-trial, error based latent updating which produces participant responding. The individual learning rate parameters within the models were estimated through hierarchical Bayesian modelling, whereby estimated group level hyperparameters influence subject level parameters. Bayesian model selection identified a best fitting model, from which individual median point estimates of learning rates were extracted and correlated to measures.

### Model specification

In the Rescorla Wagner model, an associative value, *v* was set for each CS (*v*^CS+^ and *v*^CS-^). An initial value for *v*_t=1_ was set at 0.5 for both *v*^CS^. The value was updated using the difference between the outcome, *US*, and value at each trial (prediction error), multiplied by an individual learning rate parameter (*LR*). *US* was set to 1 if the US was delivered, and 0 if not (Equation 1). Therefore, *v* was always constrained within the interval [0,1].

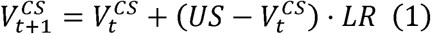

Typically in reinforcement learning models, the actions or choices tied to *v* are not ordinal. However, in this paradigm, participants select one of nine choices on an ordinal scale, and therefore a function was required to transform *v* into an ordered vector of probabilities. This was achieved through using *v* as the mean parameter of a beta distribution, with a fixed precision parameter 1 of 10. The mean and precision were re-parameterised into the two shape parameters, (a,P), of a beta distribution (Equation 2).

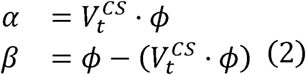

A vector, *Q*, of nine equally spaced values between zero and one inclusive, was then transformed into a vector of probabilities, *P*, through a function incorporating the beta shapes (Equation 3). This, in effect, integrated the continuous beta distribution into discrete probabilities, retaining the ordinal features of the beta distribution.

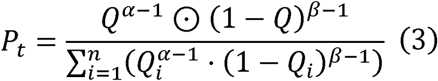

Finally, to account for stochastic deviations from the model, a lapse parameter *Lapse* was fitted. Lower values had the effect of uniformly flattening the probability vector, accounting for substantial deviations from the model by participants if required. This parameter was multiplied by *P*, and a softmax function applied to the product. This quantity was used as the theta parameter of the categorical distribution, from which participant choice of US expectancy ratings was assumed to be distributed (Equation 4).

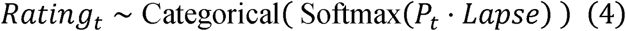

This simple, two parameter model was considered the base model (model 1a). In a combinatorial manner, additional parameters were iteratively added to this model to better account for all features of participant data. For model nomenclature within this manuscript, alterations in learning rates are represented by digits, and alterations in fitting parameters by letters, ranging from 1a to 7d.

### Fitting parameters

First, although the use of a fixed starting value is common in computational modelling, specifying a free parameter as the starting value of both *v*^CS+^ and *v*^CS-^ was tested to better incorporate participant variation in early trials (model b). Secondly, it became clear that participants generalised threat expectancy towards the CS- at the start of the extinction phase. Specifically, despite reaching the lower asymptote in the acquisition phase and responding with a US expectancy rating of one on the final CS-acquisition trial, participants would often respond significantly higher than one on the first CS-trial of extinction phase. To capture this, a free ‘jump’ parameter was estimated, and added to *v*^CS-^ prior to the extinction phase (model c). A model combining both of these fitting parameters was also fitted (model d).

### Learning rate parameters

Differing numbers and combinations of learning rates were tested to best account for participant data, and to test specifically for different learning processes occurring in the different phases, as well as in response to the different stimuli.

First, a model with two learning rates was fitted, one learning rate for CS+ trials, and one for CS-trials (model 2). This tested the hypothesis that participants learn at different rates to different cues, but that the rate is not affected by the presence of aversive stimuli (US). Second, a model with three learning rates was fitted, with a learning rate for all CS-trials, CS+ acquisition trials, and CS+ extinction trials respectively (model 3). This model specifically tests whether participants learn at different rates when acquiring and extinguishing threat, versus safety learning of a stimulus that was never threatening. Third, a model with four learning rates was tested, with a learning rate for each phase and CS. This additionally tested the CS-stimulus, which starts as ambiguous or unknown in acquisition, and is learned to be safe whereas, in extinction, no further learning should occur (model 4).

Next, learning rates were fitted which test differential learning contingent on the presence of aversive stimuli (US), which are shown to differ in those with mood or anxiety disorders (Pike & Robinson, 2022). Functionally this could only be tested within the CS+ acquisition trials, as these were the only trials to offer the delivery of the US.

First, a model with two learning rates was fitted, a learning rate for aversive trials, and a separate learning rate for non-aversive trials (model 5). This tests the hypothesis that only the presence of the US moderates the learning rate, rather than the CS. Second, a model with three learning rates was fitted, one learning rate for all CS-trials (doubling as a learning rate for non-aversive CS-trials), with a separate learning rate for aversive and non-aversive CS+ trials (model 6). This tests whether the learning rate towards non-aversive stimuli differs in threatening and non-threatening contexts. Finally, a model with five learning rates was fitted, with a learning rate for each phase and CS, save for the CS+ acquisition trials which have a separate learning rate for aversive and non-aversive trials (model 7). This final model examines all possible aspects of learning given threatening and non-threatening contexts, and aversive and non-aversive stimuli.

In summary, seven different model combinations of learning rates and four combinations of start value parameters and jump parameters were tested, making twenty-eight models in total.

### Missing data

Given the task could progress even if participants failed to enter an expectancy rating at a trial, some trial data was missing. Trials with missing data were skipped by the model, with the value update instead occurring at the next available trial with data. This prevented the need to interpolate data, or make any assumptions about learning.

### Parameter estimation

The models were fit within a hierarchical Bayesian framework. Group level hyperparameters were estimated, which constrained the subject level parameters estimated (Figure 2). Prior predictive checks were performed on all models and visually inspected to ensure hyperparameter compatibility with the underlying task structure. Uninformative priors and hyperpriors were used to allow the models to reflect the data without undue influence from the models, except for the lapse parameter which had a more informative prior (Equation 5). For learning rate, start value, and jump parameters, group level gamma hyperpriors were used to constrain the hyperparameters to positive numbers. The subject level parameters were then derived from a beta distribution, which inherently constrains the parameters within the interval [0,1]. The component shape hyperparameters were iterated by one to ensure the beta distribution did not invert. The lapse parameter, requiring a lower bound of zero, but with values near zero not desired, was derived from a gamma distribution hyperprior. The hyperparameters were also derived from gamma distributions, informed by testing, and previous literature which suggested values of around five to balance model flexibility against precision.

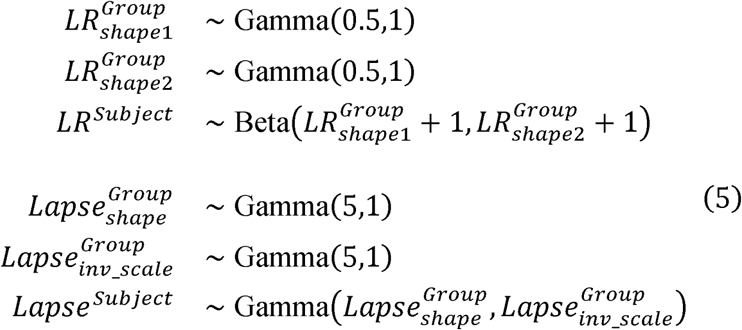

**Figure 1:**
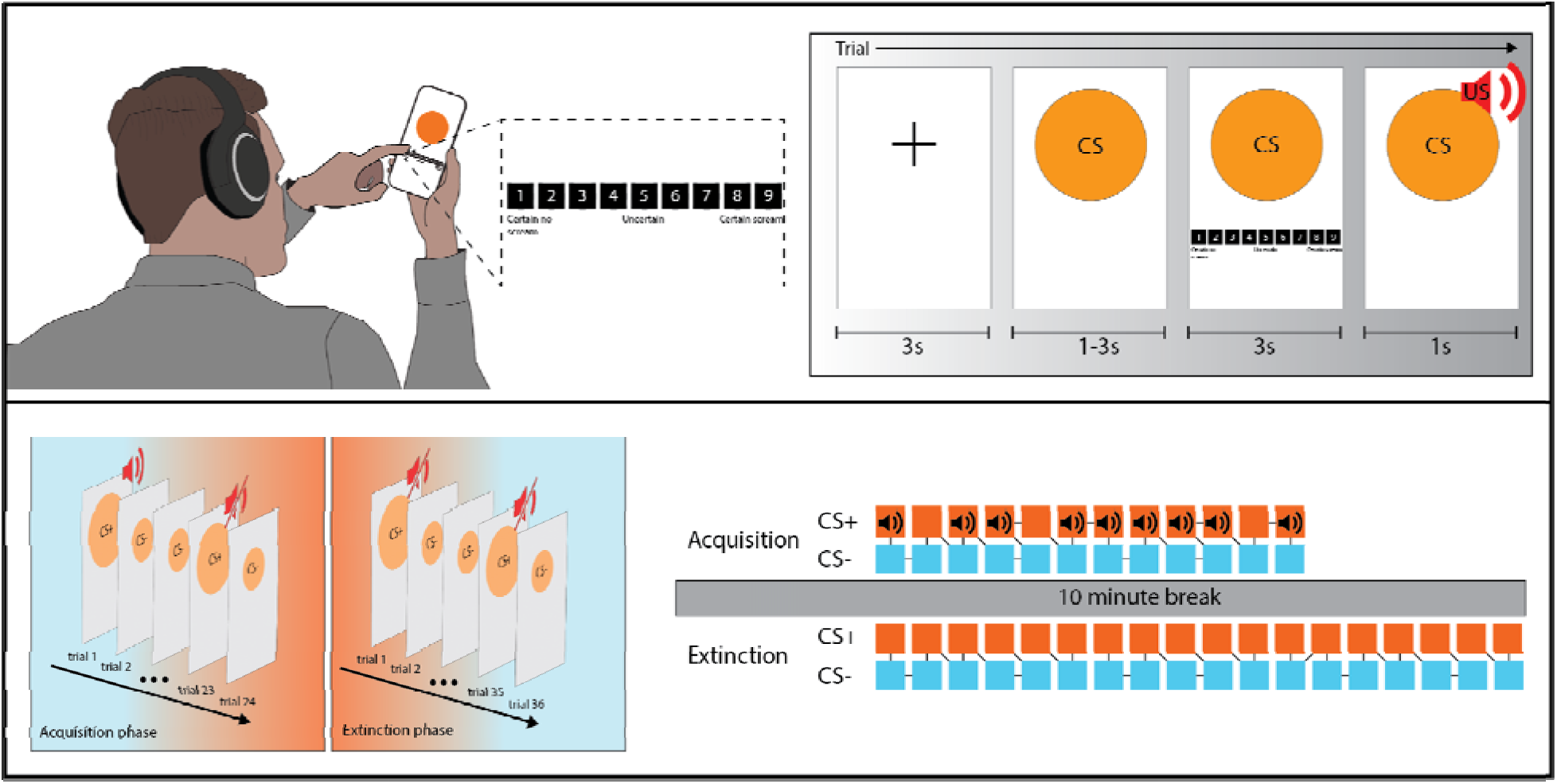
Task design. The upper panel illustrates the format of each fear conditioning trial. A fixation cross is used for the inter-trial interval. CS is presented on screen, before a US expectancy scale is presented for participants to enter their response. If the trial is reinforced with a US, this is played through participant headphones following the US expectancy rating screen. The next ITI and trial then commences. The lower panel illustrates the specific design and trial order of the paradigm in this experiment. The acquisition phase includes twenty four trials, twelve each of CS+ and CS-trials. These are presented in a pseudorandom order. CS+ is reinforced with US on 75% of occasions, i.e. nine of the twelve CS+ trials, also in a pseuodorandom order. Participants must essentially discern which CS is threatening, and which is safe. An extinction phase of thirty six trials (eighteen CS+ and eighteen CS-) follows the acquisition phase after a ten minute break. Here participants must learn that the threat CS is now safe.

**Figure 2.**
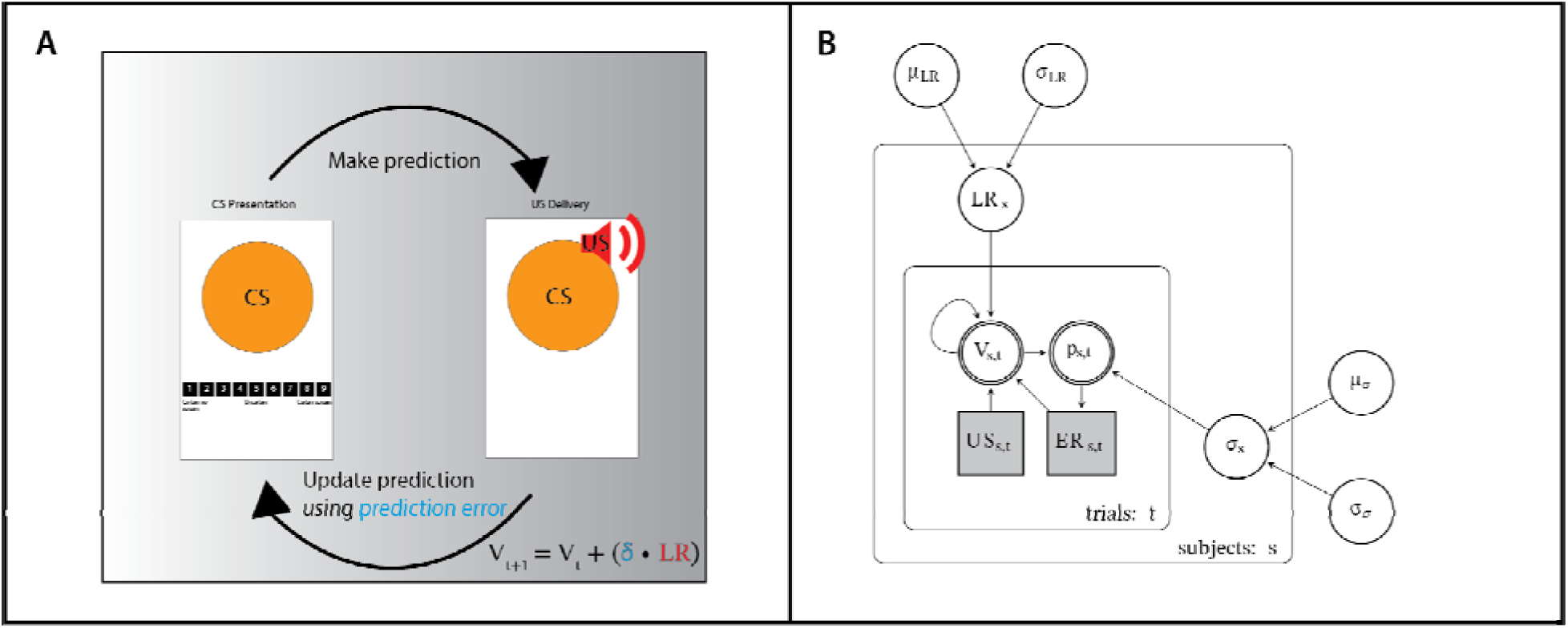
Model specification and fit A) An illustration of how the trial-by-trial structure of the fear conditioning paradigm is suited to the Rescorla Wagner model proposed. B) Hierarchical Bayesian model specification for the Rescorla Wagner model (Equation 1) applied to this task. Group level hyperpriors influence subject level parameters in the larger box. These subject level parameters determine the conditional variables (V,p) within the smaller box, representing an individual trial.

Data were simulated using fixed predefined hyperpriors to generate fixed parameters and therefore data. Each model was fitted to these data, with Markov chain Monte Carlo (MCMC) sampling used to estimate posterior probability distributions for each parameter. The median value with 95% credible intervals (95CI) of these posteriors was used as the parameter estimate, which were compared to the known input parameters. Recovery was deemed successful if the 95CI of the posterior estimate contained the true value of the parameter.

Other diagnostic checks were performed to assess chain convergence, with an R of less than 1.01 considered acceptable. Each model was then fitted to participant data using 500 warm-up iterations, and 500 sampling iterations. Following model selection, the winning model was re-run using 2000 sampling iterations to ensure maximum accuracy for parameter estimates.

### Model fit

The likelihood of the model given the trial data was calculated at each datapoint, log transformed and summed for each subject. This was performed at each phase and CS to examine the fit for each learning process modelled (Equation 6).

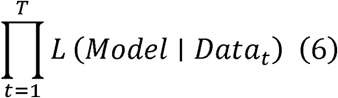

Hierarchical model fit was assessed quantitatively using the expected log pointwise predictive density (ELPD), in addition to the LOOIC and WAIC (information criteria), which penalise model complexity adding to model likelihood, and prevent overfitting. Pseudo-r^2 was also calculated, which is the ratio of model likelihood to chance (Hopkins et al., 2021).

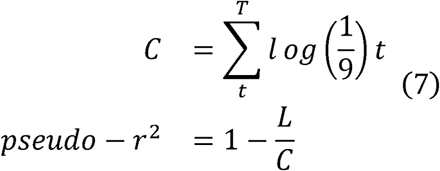

### Associations with anxiety and depression

The median point estimates of the learning rates from the winning model were correlated with GAD-7 and, separately, PHQ-8, using non-parametric Spearmans Rho correlation. This assessed the specificity of this modelling to anxiety disorders, compared to other common internalising psychiatric conditions.

### Sensitivity analysis

As part of a sensitivity analysis, a second, more stringent exclusion criteria was applied, and this cohort tested alongside the primary group. For this group, participants who reduced their headphone volume below 100%, or did not report contingency awareness, were further excluded. The modelling was applied to this cohort separately, and associations between learning rates and anxiety severity, and learning rates and depression, were tested.

### Steiger’s Z test

To compare the utility of the descriptive and computational approaches, the strength of the two sets of dependent correlations were directly compared using a modified Steiger’s Z test. The test was modified by using the absolute correlation values, as the relationship between learning rates and responding is contingent on the CS. A high learning rate in CS+ will lead to a high whole phase mean, whereas a high learning rate in CS-will lead to a low whole phase mean.

### Data and software

Task data were downloaded in csv format from the FLARe website backend, and stored on King’s College London servers. Task data were processed and manipulated into matrices using R (Version 4.4.1), for use in Stan. The models were built and specified in the Stan programming language (CmdStanR, version 2.33.1). ELPD LOO and information criteria were calculated using the loo package (version 2.6.0).

## Results

### Measures

#### Demographics

The total pooled sample (N = 235) was reduced to N = 145 following the application of post-experimental exclusion criteria. There was a significant difference observed in the median GAD-7 score between the pooled study groups (H=7.17, p=0.028). However, this difference did not survive the post-experiment exclusion criteria. No other significant differences in demographics or outcome measures were noted between the pooled study groups. Likewise, no significant differences were noted between groups before and after the experiment exclusion criteria were applied (Table 1).

### Task

#### Primary Analysis

##### Associations of descriptive measures with anxiety severity

In keeping with our pre-registered hypotheses, end phase CS discrimination scores, the difference between final CS+ and CS-expectancy rating for each phase, were correlated with GAD-7 scores (Table 2). As predicted, both acquisition and extinction phase CS discrimination scores were significantly associated with anxiety severity (r = −0.179, p = 0.032 & r = 0.185, p = 0.026 respectively). Similarly, significant associations were observed between all whole phase means (the mean expectancy rating towards each CS across each phase) and GAD-7, except the acquisition phase CS+ mean rating.

**Table 2.**
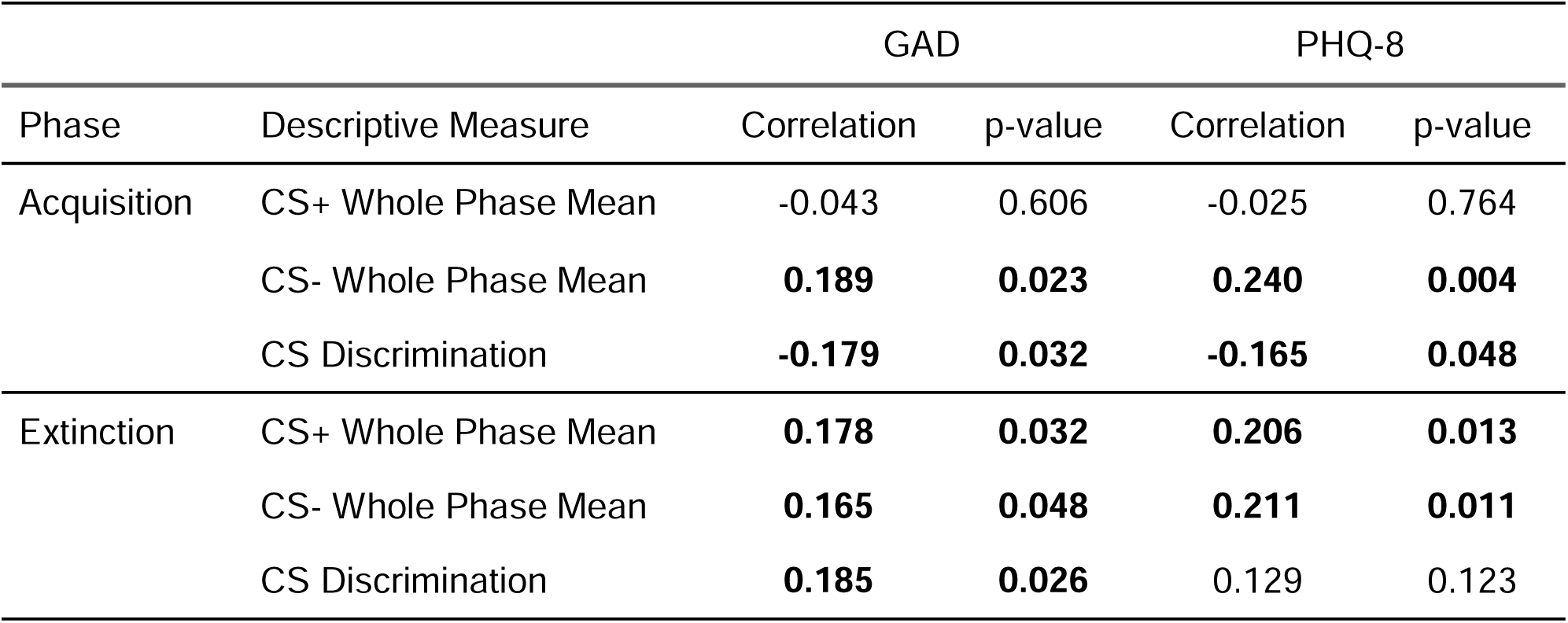
Correlations of fear conditioning descriptive measures with anxiety severity, and depression (n = 145)

### Computational modelling

#### MCMC computational checks

All models tested achieved post-warmup chain convergence, with no model displaying a Gelman-Rubin statistic (*R*) above 1.01, nor were any divergences discovered. Visual checks of hyperparameter traceplots and rank plots supported this quantitative assessment of convergence (Baribault & Collins, 2023). Posterior predictive checks were performed to assess model fit, with the model generating data which closely matched the observed data upon visual inspection (Figure 3, panel A). This was corroborated quantitatively, with the whole phase mean of generated data compared to whole phase means of participant data through correlation (Figure 3, panel C). This demonstrated that, on average, individual differences in generated data matched that of real data, and that the model is capturing observed behaviour.

**Figure 3.**
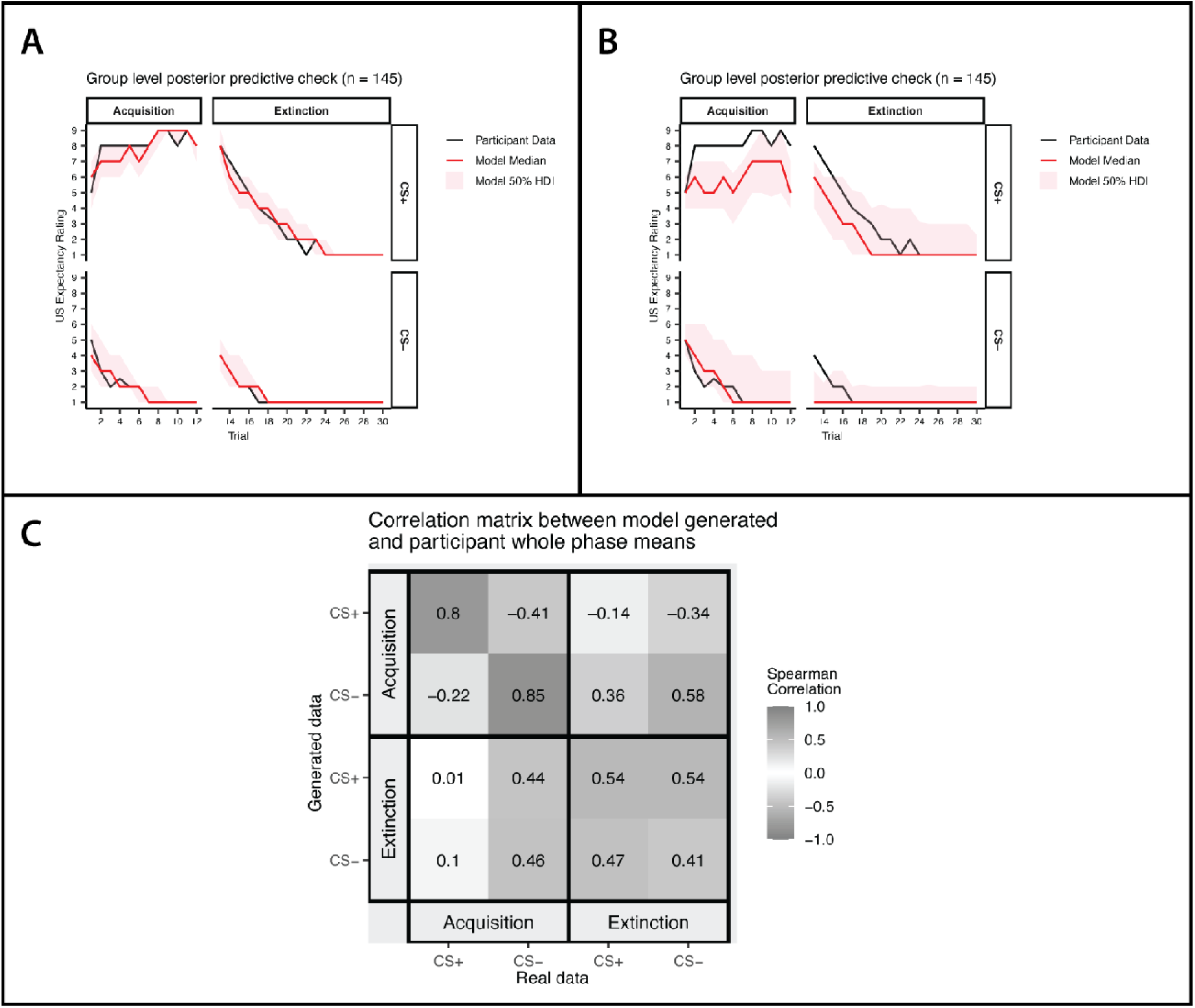
Posterior predictive check A) Group level posterior predictive check (PPC) comparing model generated data from the winning five-learning rate model, to real participant data. The shaded area represents the 50% highest density interval (HDI), with the solid red line indicating the median generated expectancy rating. B) By comparison, a group level PPC for the single learning rate model, demonstrating a much poorer fit and wider HDI. C) A correlation matrix of generated whole phase means (y axis) to real whole phase means (x axis). The left to right diagonal compares like with like, indicating a better fit in the acquisition phase compared to extinction.

#### Model comparison and selection

The model with the overall highest likelihood given the data was model 7d, which contained five learning rate parameters, free start value parameters, and the CS-extinction phase jump parameter (Supplementary Information). This held when examining the fits at the level of each phase and CS, except for extinction phase CS-, where model 4d, the four learning rates model, was best fitting. However, the difference in ELPD LOO between these two models did not exceed five times the standard error, therefore model 7d was used for subsequent analysis to ensure consistency and comparability across phases.

### Associations of computational parameters with anxiety severity

#### Learning rates

As predicted, the threat extinction (CS+ during extinction) and safety learning (CS-during acquisition) rates were both negatively associated with anxiety severity, such that as anxiety severity increases, the rates of extinction and safety learning decrease (r = −0.218, p = 0.008 & r = −0.214, p = 0.01 respectively). In contrast, and against prediction, the rate of threat acquisition was not associated with anxiety severity (r = −0.07, p = 0.404). The rate of threat extinction was also negatively associated with depression severity (r = −0.226, p = 0.006) (Table 3).

**Table 3.** Learning Rate Correlations with Anxiety Severity.

| Spearman correlations between model parameters and measures |  |  |  |  |
| --- | --- | --- | --- | --- |
| Parameter | GAD-7 |  | PHQ-8 |  |
|  | Medium<br>(n=145) | Strict<br>(n=88) | Medium<br>(n=145) | Strict<br>(n=88) |
| Acquisition CS+ Learning Rate (US+) | -0.070 (p=0.404) | -0.155 (p=0.150) | -0.035 (p=0.678) | -0.006 (p=0.953) |
| Acquisition CS+ Learning Rate (US-) | -0.097 (p=0.247) | **-0.299 (p=0.005)** | -0.057 (p=0.492) | -0.198 (p=0.065) |
| Acquisition CS- Learning Rate | **-0.218 (p=0.008)** | **-0.317 (p=0.003)** | -0.136 (p=0.102) | -0.140 (p=0.192) |
| Extinction CS+ Learning Rate | **-0.214 (p=0.010)** | **-0.334 (p=0.001)** | **-0.226 (p=0.006)** | **-0.256 (p=0.016)** |
| Extinction CS- Learning Rate | -0.069 (p=0.413) | -0.147 (p=0.172) | -0.099 (p=0.237) | -0.134 (p=0.212) |
| Lapse Rate | -0.057 (p=0.498) | -0.132 (p=0.220) | -0.156 (p=0.061) | **-0.215 (p=0.044)** |
| CS+ Initial Value | 0.047 (p=0.572) | 0.079 (p=0.465) | 0.038 (p=0.648) | 0.018 (p=0.865) |
| CS- Initial Value | 0.039 (p=0.642) | -0.074 (p=0.493) | 0.126 (p=0.131) | 0.002 (p=0.988) |
| Extinction CS- Jump Value | -0.059 (p=0.483) | -0.076 (p=0.481) | -0.067 (p=0.426) | -0.069 (p=0.526) |

### Sensitivity analysis

The total sample was reduced to n = 88 following the application of strict exclusion criteria indicating task inattentiveness or non-compliance. No significant differences were noted in outcome measures between the groups before and after the exclusion criteria were applied (Table 1).

In this sample, the associations between the threat extinction and safety learning rates and anxiety severity were stronger and remained significant (r = −0.317, p=0.003 & r = −0.334, p=0.001). However, in addition, the learning rate towards unreinforced trials in acquisition phase CS+ trials became negatively associated with anxiety severity (r = −0.299, p=0.005).

### Steigers Z comparison of correlations

Although the relationship between computational parameters and anxiety severity was uniformly higher than the corresponding descriptive measure associations, these differences were not significant where direct comparisons were possible (Table 4).

**Table 4.** Steiger’s Z Test Results.

| Stan Parameter | WPM Measure | T Statistic | p-value |
| --- | --- | --- | --- |
| Acquisition CS+ Learning Rate (US+) | Acquisition CS+ WPM | 0.330 | 0.742 |
| Acquisition CS+ Learning Rate (US-) | Acquisition CS+ WPM | 0.482 | 0.630 |
| Acquisition CS- Learning Rate | Acquisition CS- WPM | 0.349 | 0.727 |
| Extinction CS+ Learning Rate | Extinction CS+ WPM | 0.873 | 0.384 |
| Extinction CS- Learning Rate | Extinction CS- WPM | -1.238 | 0.218 |

## Discussion

This study demonstrates that distinct learning rates can be modelled for participants undergoing smartphone delivered fear conditioning, and that these learning rates demonstrate associations with anxiety severity. Specifically, in support of existing literature, safety learning, and threat extinction learning were negatively associated with anxiety severity, implying that those with greater anxiety struggle to learn that objects or situations are safe (Abend et al., 2022; Duits et al., 2015). Conversely, despite an association between CS discrimination and anxiety severity in the acquisition phase, threat acquisition learning rates were not associated with anxiety severity. This may indicate that safety learning is more relevant to the experience of anxiety than threat learning.

The computational model offered a reasonable fit to the data, and uniformly accounted for more of the evidence numerically than descriptive approaches, albeit not significantly where comparisons were possible. This is likely an effect of the simple structure of the task, where the simple asymptotic curves are well accounted for by a descriptive measure. Interestingly, in support of this, model fit was superior in the more volatile acquisition phase, than in the more predictable extinction phase, demonstrated through higher correlations between generated and participant data and lower mean squared error (Figure 3, panel C).

However, a strong advantage of this computational modelling, is a reduction of the analytic flexibility seen with use of descriptive statistics (Lonsdorf et al., 2022). Here, a principled approach of model fitting and selection was used, applied to all trial data, producing a single model capable of explaining the causal effects which can only be indirectly measured through the many descriptive statistics seen in the literature. Whilst computational modelling is essential in more complex task structures, this study highlights its value in simpler tasks, offering a clear analytic rationale.

One aim within the field of computational psychiatry is to unearth previously hidden differences between psychiatric conditions (Friston et al., 2017). Depression is often comorbid with generalised anxiety disorder, yet they are conceived as having separate underlying mechanisms. Specifically, the disordered threat processing of anxiety is not usually included in theoretical models of the aetiology of depression. In this study, a supplementary analysis of the associations between a depression scale (PHQ-8) and learning rates, showed no difference from the primary analysis of those between GAD-7 and learning rates (Table 3). This may indicate a common mechanism to both, or that the task is not sensitive to the differences between the two symptom severity measures, which are themselves highly correlated.

Interestingly, applying more stringent exclusion criteria, whilst reducing the sample size substantially, demonstrated not only stronger associations between learning rates and GAD-7, but eliminated the associations to PHQ-8, perhaps offering greater diagnostic specificity from this approach (Table 3).

### Limitations

Remotely collected data inherently lacks experimenter control, and therefore adherence to the task cannot be guaranteed. This is likely compounded by the repetitive and aversive nature of the task. These biases are partially offset through questioning the subjects following the task, including questions about headphone removal or contingency awareness. There is no guarantee these questions are answered honestly, so the biases cannot be completely discounted. Only a third of participants engaged with the task as fully intended, surviving the strictest exclusion criteria. Indeed, model fit, and associations with anxiety, were strengthened when maximally controlling for deviations from the task, indicating that these control measures offset some of the disadvantages posed by the limitations of remote collection. The large sample sizes collected through FLARe further offset the loss of sample size through the application of exclusion criteria.

The GAD-7 scale is predominantly a measure of generalised anxiety disorder, whereas fear conditioning is shown to be more relevant to fear-based disorders, including specific phobias and PTSD (Lonsdorf & Merz, 2017). These disorders tend to have a specific threat at their core (analogous to a specific CS), in contrast to the more abstract or distal threats perceived in generalised anxiety disorder. Therefore, we might expect to see even stronger associations towards a measure of fear-based disorders.

The standard Rescorla Wagner model is sufficiently parameterised for estimating learning rates, though the accuracy of them could be questioned, on technical and theoretical grounds. We also attempted to fit other, more complex models, containing more parameters to account for further learning processes. However, these models universally proved to be computationally intractable, or failed to converge chains, thus not offering reasonable parameter estimates (Baribault & Collins, 2023). These issues likely stem from, and are compounded by, the relative lack of trial-by-trial data available for each phase, compared to analogous modelling tasks in the literature using hundreds of trials (Pike & Robinson, 2022).

Whilst reinforcement learning is a reasonable theoretical model to apply to this data, and almost certainly accounts for some of the behaviour observed, other cognitive and learning processes are likely to co-occur during the task (Lonsdorf et al., 2017b). The lapse parameter attempts to account for all these overlapping processes, in addition to measurement error.

However other models could better account for this ‘deviance’ from the Rescorla Wagner model, including counter-factual reasoning and reversal learning models (Gershman & Hartley, 2015; Zika et al., 2023). Similar studies suggest probabilistic models may better account for fear conditioning, albeit using physiological conditioned responses with far greater response noise and measurement error (Tzovara et al., 2018).

### Future work

Future work may look to add complexity to the task structure, generating data in which models would have stronger predictive power over descriptive measures. In addition, confirmatory studies using the more stringent exclusion criteria, to counterbalance the lack of experimenter control inherent to remotely delivered paradigms, may further strengthen the associations elicited through computational modelling.

This study cannot ascribe a direction to the mechanism demonstrated. It is reasonable to suggest that those with a pre-existing slow learning rate may be at risk of developing anxiety disorders, but equally it could be the case that those with anxiety disorders then develop a slow learning rate as a symptom, contributing to the persistence of the disorders. Future work might look to repeat this novel approach longitudinally to establish a causal direction.

### Summary

Computational modelling offers a credible account of the learning processes contributing to anxiety severity, over and above traditional descriptive measures which can only hint at them. This theory driven, reproducible analytic method paves the way for larger scale studies using similar computational modelling in smartphone delivered paradigms, to further instigate these novel mechanistic insights into anxiety.

