## Supplementary figures and tables for "Computational modelling reveals slower safety learning and threat extinction are associated with higher anxiety severity in remote fear conditioning"

Supplementary Information

### Prior predictive checks

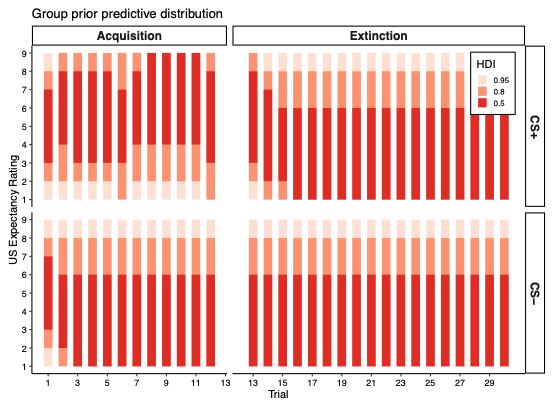

Prior predictive check for model with single learning rate and no extra fitting parameters (model 1a). This fails to provide model space for participant behaviour in some sections

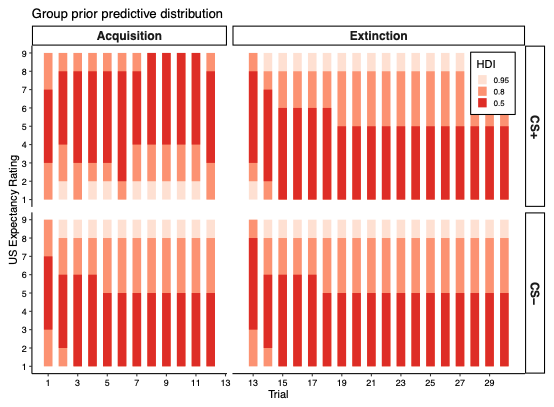

Prior predictive check for the winning model, with five learning rate parameters and three fitting parameters (model 7d). This provides model space for participant data to be modelled accurately.

### Model comparison tables

#### All phases combined

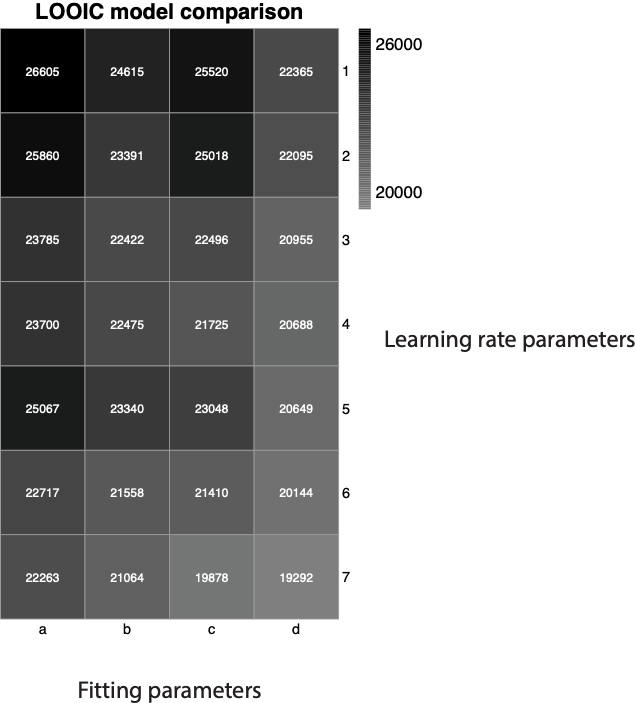

LOOIC model comparison matrix for all 28 models tested

|  | | **Parameters** | | | | **Metrics** | | | | | |
| --- | --- | --- | --- | --- | --- | --- | --- | --- | --- | --- | --- |
| **Model** | **Name** | **Total** | **LR** | **Start** | **Jump** | **Log-likelihood** | **LOOIC** | **WAIC** | **BIC** | **Pseudo R²** | **LOO** |
| 1c | lr1_single_fixed_jump | 3 | 1 | 0 | 1 | -2399215.474 | 25520.282 | 25971.31 | 64.614 | 0.361 | -12760.141 |
| 1a | lr1_single_fixed_nojump | 2 | 1 | 0 | 0 | -2533454.26 | 26605.494 | 27164.607 | ****59.179**** | 0.325 | -13302.747 |
| 1d | lr1_single_free_jump | 5 | 1 | 2 | 1 | -2099620.565 | 22365.419 | 23666.74 | 74.947 | 0.441 | -11182.709 |
| 1b | lr1_single_free_nojump | 4 | 1 | 2 | 0 | -2296692.742 | 24614.974 | 25852.886 | 70.595 | 0.388 | -12307.487 |
| 2c | lr2_cspc_fixed_jump | 4 | 2 | 0 | 1 | -2313063.62 | 25017.735 | 26390.698 | 70.878 | 0.384 | -12508.868 |
| 2a | lr2_cspc_fixed_nojump | 3 | 2 | 0 | 0 | -2425821.631 | 25860.478 | 27191.231 | 65.072 | 0.354 | -12930.239 |
| 2d | lr2_cspc_free_jump | 6 | 2 | 2 | 1 | -2049759.947 | 22094.503 | 23211.88 | 81.837 | 0.454 | -11047.251 |
| 2b | lr2_cspc_free_nojump | 5 | 2 | 2 | 0 | -2180016.38 | 23391.105 | 24422.112 | 76.333 | 0.419 | -11695.552 |
| 3c | lr2_posneg_fixed_jump | 4 | 2 | 0 | 1 | -2157918.566 | 23047.707 | 23263.478 | 68.203 | 0.425 | -11523.854 |
| 3a | lr2_posneg_fixed_nojump | 3 | 2 | 0 | 0 | -2340527.309 | 25067.086 | 25792.111 | 63.602 | 0.377 | -12533.543 |
| 3d | lr2_posneg_free_jump | 6 | 2 | 2 | 1 | -1929506.71 | 20649.237 | 20792.125 | 79.763 | 0.486 | -10324.618 |
| 3b | lr2_posneg_free_nojump | 5 | 2 | 2 | 0 | -2152127.266 | 23339.664 | 24578.077 | 75.852 | 0.427 | -11669.832 |
| 4c | lr3_cspacqext_csm_fixed_jump | 5 | 3 | 0 | 1 | -2114543.962 | 22495.571 | 22979.959 | 75.204 | 0.437 | -11247.786 |
| 4a | lr3_cspacqext_csm_fixed_nojump | 4 | 3 | 0 | 0 | -2253001.888 | 23784.888 | 24669.178 | 69.842 | 0.4 | -11892.444 |
| 4d | lr3_cspacqext_csm_free_jump | 7 | 3 | 2 | 1 | -1950932.94 | 20954.9 | 21407.935 | 87.882 | 0.48 | -10477.45 |
| 4b | lr3_cspacqext_csm_free_nojump | 6 | 3 | 2 | 0 | -2093060.269 | 22421.722 | 22733.131 | 82.583 | 0.443 | -11210.861 |
| 5c | lr3_cspposneq_csm_fixed_jump | 5 | 3 | 0 | 1 | -2008271.16 | 21409.815 | 21573.186 | 73.372 | 0.465 | -10704.908 |
| 5a | lr3_cspposneq_csm_fixed_nojump | 4 | 3 | 0 | 0 | -2149653.845 | 22717.221 | 22839.474 | 68.06 | 0.428 | -11358.61 |
| 5d | lr3_cspposneq_csm_free_jump | 7 | 3 | 2 | 1 | -1880168.452 | 20144.119 | 20170.681 | 86.662 | 0.499 | -10072.059 |
| 5b | lr3_cspposneq_csm_free_nojump | 6 | 3 | 2 | 0 | -2014910.468 | 21558.293 | 21627.165 | 81.236 | 0.463 | -10779.146 |
| 6c | lr4_cspacqext_csmaqext_fixed_nojump | 5 | 4 | 0 | 0 | -2228597.618 | 23699.645 | 24454.166 | 77.171 | 0.407 | -11849.822 |
| 6a | lr4_cspacqext_csmaqext_free_nojump | 7 | 4 | 2 | 0 | -2080351.246 | 22474.557 | 23048.15 | 90.113 | 0.446 | -11237.278 |
| 6d | lr4_cspaqext_csmaqext_fixed_jump | 6 | 4 | 0 | 1 | -2048224.5 | 21725.327 | 22243.526 | 81.81 | 0.455 | -10862.664 |
| 6b | lr4_cspaqext_csmaqext_free_jump | 8 | 4 | 2 | 1 | -1923890.766 | 20687.778 | 20984.372 | 95.165 | 0.488 | -10343.889 |
| 7c | lr5_cspaqposneg_cspext_csmaqext_fixed_jump | 7 | 5 | 0 | 1 | -1865452.841 | 19877.804 | 20223.854 | 86.408 | 0.503 | -9938.902 |
| 7a | lr5_cspaqposneg_cspext_csmaqext_fixed_nojump | 6 | 5 | 0 | 0 | -2069167.634 | 22263.333 | 22965.953 | 82.171 | 0.449 | -11131.666 |
| 7d | lr5_cspaqposneg_cspext_csmaqext_free_jump | 9 | 5 | 2 | 1 | ****-1787621.401**** | ****19291.665**** | ****19654.715**** | 100.565 | ****0.524**** | ****-9645.832**** |
| 7b | lr5_cspaqposneg_cspext_csmaqext_free_nojump | 8 | 5 | 2 | 0 | -1946379.092 | 21064.231 | 21231.792 | 95.553 | 0.482 | -10532.116 |

### Root mean square error

| phase | rmse |
| --- | --- |
| acquisition_CS+ | 0.6888018 |
| acquisition_CS- | 0.6930095 |
| extinction_CS+ | 1.8026228 |
| extinction_CS- | 2.7877411 |

Root mean square error between real and model generated whole phase means
